# Utilization of a *Histoplasma capsulatum* zinc reporter reveals the complexities of fungal sensing of metal deprivation

**DOI:** 10.1101/2023.11.14.567133

**Authors:** Logan T. Blancett, Heather M. Evans, Kathleen Candor, William R. Buesing, Julio A. Landero Figueroa, George S. Deepe

## Abstract

*Histoplasma capsulatum* is a dimorphic fungal pathogen acquired via inhalation of soil-resident spores. Upon exposure to mammalian body temperatures, these fungal elements transform into yeasts that reside primarily within phagocytes. Macrophages (MΦ) provide a permissive environment for fungal replication until T cell-dependent immunity is engaged. MΦ activated by granulocyte-MΦ colony stimulating factor (GM-CSF) induce metallothioneins (MTs) that bind zinc (Zn) and deprive yeast cells of labile Zn, thereby disabling fungal growth. Prior work demonstrated that the high affinity zinc importer, ZRT2, was important for fungal survival in vivo. Hence, we constructed a yeast cell reporter strain that expresses green fluorescent protein (GFP) under the control of this importer. This reporter accurately responds to medium devoid of Zn. ZRT2 expression increased (∼5-fold) in GM-CSF, but not interferon-γ, stimulated MΦ. To examine the in vivo response, we infected mice with reporter yeasts and assessed ZRT2 expression at 0-, 3-, 7-, and 14-days post-infection (dpi). ZRT2 expression minimally increased at 3-dpi and peaked on 7-dpi, corresponding with onset of adaptive immunity. We discovered that the major phagocyte populations that restrict Zn to the fungus are interstitial MΦ and exudate MΦ. Neutralizing GM-CSF blunted control of infection but unexpectedly increased ZRT2 expression. This increase was dependent on another cytokine that activates MΦ to control *H. capsulatum* replication, M-CSF. These findings illustrate the reporter’s ability to sense Zn *in vitro* and *in vivo* and correlate ZRT2 activity with GM-CSF and M-CSF activation of MΦ.

**Importance:** Phagocytes use an arsenal of defenses to control replication of *Histoplasma* yeasts, one of which is limitation of trace metals. On the other hand, *H. capsulatum* combats metal restriction by upregulating metal importers such as the Zn importer ZRT2. This transporter contributes to *H. capsulatum* pathogenesis upon activation of adaptive immunity. We constructed a fluorescent ZRT2 reporter to probe *H. capsulatum* Zn sensing during infection and exposed a role for M-CSF activation of macrophages when GM-CSF is absent. These data highlight the ways in which fungal pathogens sense metal deprivation in vivo and reveal the potential of metal-sensing reporters. The work adds a new dimension to studying how intracellular pathogens sense and respond to the changing environments of the host.

## Introduction

The ability of microbes to acquire nutrients such as essential metals from the host is paramount for successful colonization and proliferation. To control infection, host cells must restrict access to metals via a process termed “nutritional immunity” (1). Originally described as iron (Fe) restriction or toxicity, the definition now extends to control of zinc (Zn) and copper (Cu) (2–5). Mammalian hosts regulate extracellular and intracellular Zn concentrations by employing Zn-binding proteins including calprotectin or metallothioneins (MTs). The latter calibrate intracellular Zn concentrations by binding and shuttling Zn into cellular organelles (4, 6, 7). Thus, microbe and host battle for Zn to survive (8–14).

*H. capsulatum* is a thermally dimorphic fungus with a worldwide distribution and is endemic to the US (15, 16). Control of infection depends on cooperativity among T cells, macrophages (MΦ), and dendritic cells (DCs), but largely relies on stimulating the antifungal machinery of MΦ (17). Interferon (IFN)γ and granulocyte-monocyte colony stimulating factor (GM-CSF) are two cytokines that arm these phagocytes to limit intracellular growth of yeasts by constraining access to trace metals (18). IFNγ restricts Cu and Fe (19–21), and GM-CSF restricts Zn (22).

Deprivation of Zn in GM-CSF-activated MΦ is accomplished by MTs 1&2-mediated sequestration. Consequently, yeast cells are starved of Zn, rendering them more susceptible to killing by MΦ-induced reactive oxygen species (ROS) (22–24). One mechanism by which *Histoplasma* copes with Zn limitation is upregulating the high affinity Zn transporter *ZRT2*. This gene is crucial for optimal survival of the fungus in mice, especially during the genesis of cellular immunity, ∼7-days post-infection (dpi) (25).

While the GM-CSF-dependent pathway by which MΦ restrict Zn has been examined, no studies exist probing Zn availability in lung MΦ populations or how the fungus senses changes in this metal. To accomplish this, we constructed a reporter strain expressing green fluorescent protein (GFP) under the control of this Zn importer (ZRT2-GFP). This reporter accurately responded to medium devoid of Zn and expression of ZRT2-GFP increased in GM-CSF-, but not IFNγ, -activated MΦ. In vivo, ZRT2-GFP expression increased concomitant with the onset of adaptive immunity. Disruption of GM-CSF signaling in vivo blunted host defenses but unexpectedly provoked heightened ZRT2-GFP expression. Increased reporter expression in GM-CSF-deficient mice depended on M-CSF, another cytokine that activates MΦ to inhibit growth of *H. capsulatum* (26). This study highlights the complex nature of Zn regulation in *Histoplasma*-infected MΦ and demonstrates that sensing Zn deprivation is uncoupled from host restriction of intracellular replication.

## Results

### Fidelity of the ZRT2-GFP Zn reporter

The *H. capsulatum* genome encodes four putative Zn transporters, designated ZRT1, ZRT2, ZRT3, and ZRC1, and a Zn-dependent transcriptional regulator ZAP1. We analyzed transcriptional changes of Zn-responsive genes in standard (4 μM), low (0.1 µM) and high Zn (100 µM) medium. In the former, transcription of *ZRT2*, *ZRT3*, and *ZAP1* was increased, *ZRC1* decreased, and *ZRT1* was unchanged (Fig. 1a). ZRT2 is a major importer of Zn and required for optimal growth of *Histoplasma* in mouse lungs (25). We hypothesized that creation of a ZRT2 reporter strain would facilitate detection of Zn availability to yeasts in host cell phagosomes. To this end, we fused the gene encoding green-fluorescent protein (GFP) to the *ZRT2* promoter (ZRT2-GFP). The construct was inserted into *H. capsulatum* yeasts that constitutively express tdTomato (TOM) to enable simultaneous detection of phagocyte infection and phagosomal Zn availability (Fig. 1b). The potential outcomes of this reporter in a low and high Zn environment are depicted in Fig. 1b (right hand panel). The reporter strain grew equally well as wild-type yeast cells in liquid culture and in mouse lungs (Fig. S1). Zn reporter yeasts exhibited a 4-fold higher expression in low Zn compared to standard and high Zn medium (Fig. S2), and a 6-fold increase in medium containing 10 µM of the Zn chelator *N*,*N*,*N′*,*N′*-tetrakis(2-pyridinylmethyl)-1,2-ethanedimine (TPEN) (Fig. S2). An isogenic strain of *H. capsulatum* that expresses GFP under control of the constitutive *Translation Elongation Factor 1 (TEF1)ɑ* promoter did not exhibit increased GFP (Fig. S2). Thus, the ZRT2-GFP reporter senses changes in Zn availability.

**Figure 1.**
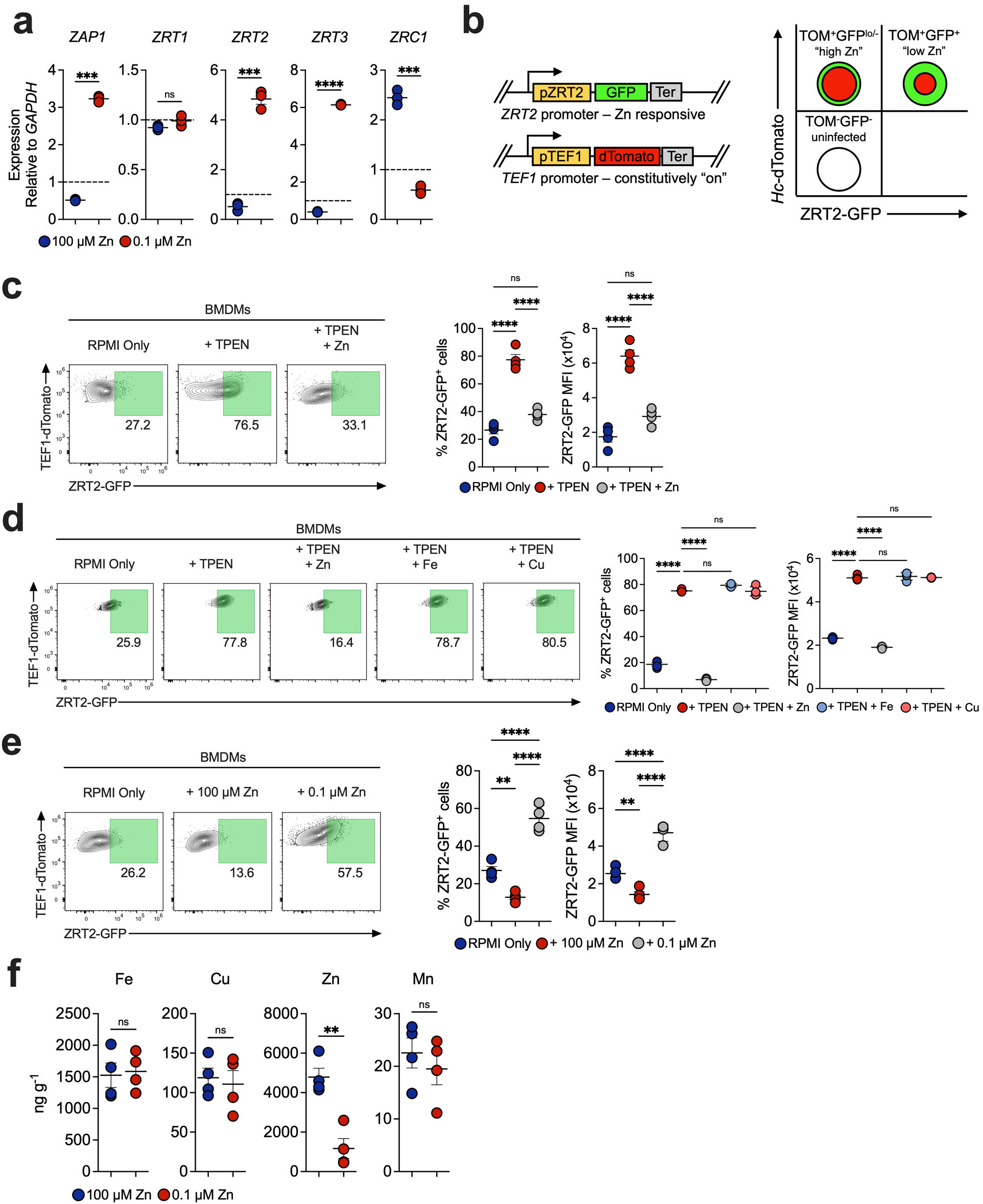
Fidelity of the ZRT2-GFP Zn Reporter. (**a**) qRT-PCR of *Histoplasma* yeasts incubated for 4 hours in high Zn (100 µM) or low Zn (0.1 µM). Expression normalized to yeast cells in HMM as represented by dashed line. (**b**) Schematic of the ZRT2-GFP Zn reporter construction. Right hand figure illustrates expected results with flow cytometric analysis of infected and uninfected cells. ZRT2-GFP is constitutively expressed therefore there will always be detection of GFP. **(c)** ZRT2-GFP % and GFP MFI from unstimulated, 48 hpi ZRT2-GFP-infected BMDMs incubated in RPMI only, RPMI + 100 µM Zn, or RPMI + 0.1 µM Zn. **(d)** RPMI only, RPMI + 10 µM TPEN, or RPMI + TPEN + 100 µM Zn.**(e)** ZRT2-GFP in medium with low or high Zn. (**f**) ICP-MS derived metal concentrations from *Histoplasma* yeasts extracted from 48-hpi ZRT2-GFP-infected BMDMs incubated in high Zn (100 µM) or low Zn (0.1 µM).

To demonstrate that the reporter responds to changes in Zn availability during intracellular residence, we cultured reporter-infected, bone marrow-derived MΦ (BMDMs) in RPMI (2 µM Zn) with or without TPEN and measured ZRT2-GFP expression 48-hours post-infection (hpi). TPEN increased the proportion of ZRT2-GFP^+^ BMDMs by 3-fold and GFP median fluorescence intensity (MFI) by 2-fold (Fig. 1c). Since TPEN chelates other divalent cations, we examined the specificity for Zn (27). Only Zn, but not Cu or Fe, diminished ZRT2 expression (Fig. 1d). We also incubated Zn reporter-infected BMDMs in Zn-free medium supplemented with high (100 µM) or low Zn (0.1 µM). The Zn reporter manifested a 4-fold elevation in ZRT2-GFP^+^ BMDMs and increased GFP MFI in low Zn (Fig. 1e). To confirm that Zn was the sole divalent cation responsible for the changes, we analyzed metal content in yeast cells by inductively coupled plasma-mass spectrometry (ICP-MS): only Zn was diminished (Fig. 1f).

### GM-CSF stimulation of BMDMs Induces ZRT2

GM-CSF stimulation of MΦ enhances expression of MTs 1 & 2 that sequester Zn from yeasts (22). To determine if the ZRT2 reporter responds to GM-CSF-induced Zn deprivation, infected BMDMs were exposed to vehicle, GM-CSF, or IFNγ, and at 48-hpi, cells were analyzed by flow cytometry. While no changes in ZRT2-GFP were observed in vehicle or IFNγ-exposed cells (Fig. 2a), GM-CSF induced a 50% increase in ZRT2-GFP^+^ BMDMs and a 3-fold increase in GFP MFI (Fig. 2b). The addition of 100 µM of Zn to GM-CSF-stimulated BMDMs decreased the proportion of ZRT2-GFP^+^ cells and GFP MFI. Zn deprivation enhances ROS production in MΦ (28); therefore, we exposed resting and GM-CSF-stimulated BMDMs to the phagosomal NADPH oxidase inhibitor apocynin (APO) (29). Inhibition of ROS did not affect ZRT2-GFP expression (Fig. 2c). The Zn reporter senses a low Zn environment in phagosomes of GM-CSF-activated MΦ. The data reveal that GM-CSF, not IFNγ, provokes a Zn-deficient phagosomal environment.

**Figure 2.**
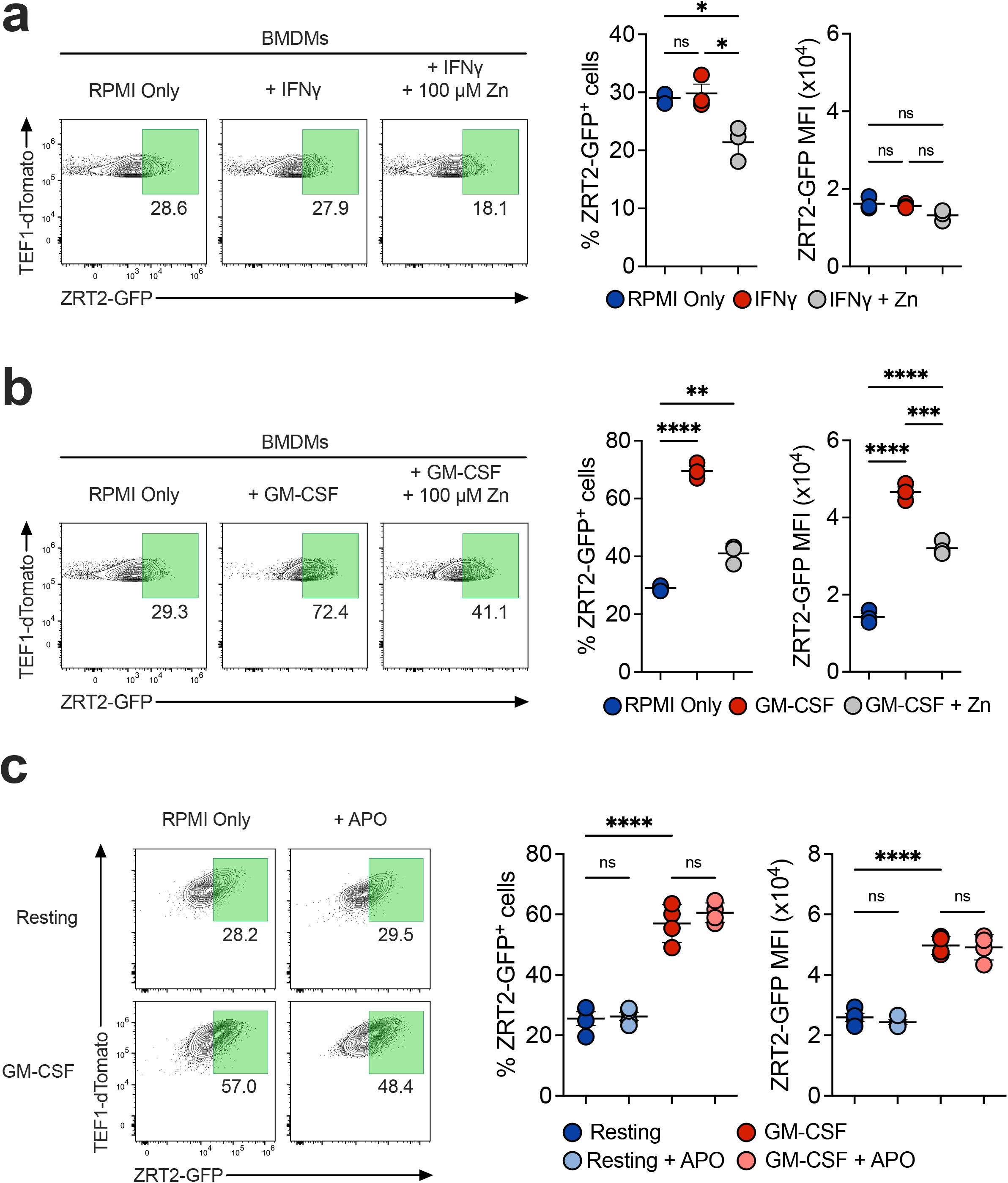
GM-CSF Stimulation Induces ZRT2. ZRT2-GFP-infected BMDMs stimulated with (**a**) 10 ng/mL IFNγ or (**b**) GM-CSF for 48 hours. (**c**) resting or GM-CSF stimulated BMDMs treated with 200 µM Apocynin or vehicle (DMSO) for 48 hours.

### ZRT2-GFP expression in lung cell populations

To better understand the in vivo dynamics of Zn deprivation during infection, we challenged WT mice with reporter yeasts and analyzed ZRT2-GFP in total leukocytes (CD45^+^) at 3-, 7-, and 14-dpi, corresponding to onset (innate), peak (adaptive), and resolution of infection, respectively (Fig. 3a) (17). ZRT2 is consistently detected in yeasts even in Zn replete conditions. Thus, we needed to establish a threshold to distinguish ZRT2 “basal” (ZRT2-GFP^-^) vs. “induced” (ZRT2-GFP^+^). Accordingly, we infected mice for 2 hours and analyzed ZRT2-GFP in lung leukocytes. To be stringent, we established that an MFI 4 SD above the threshold will be referred to as ZRT2-GFP^+^ (Fig. S3).

**Figure 3.**
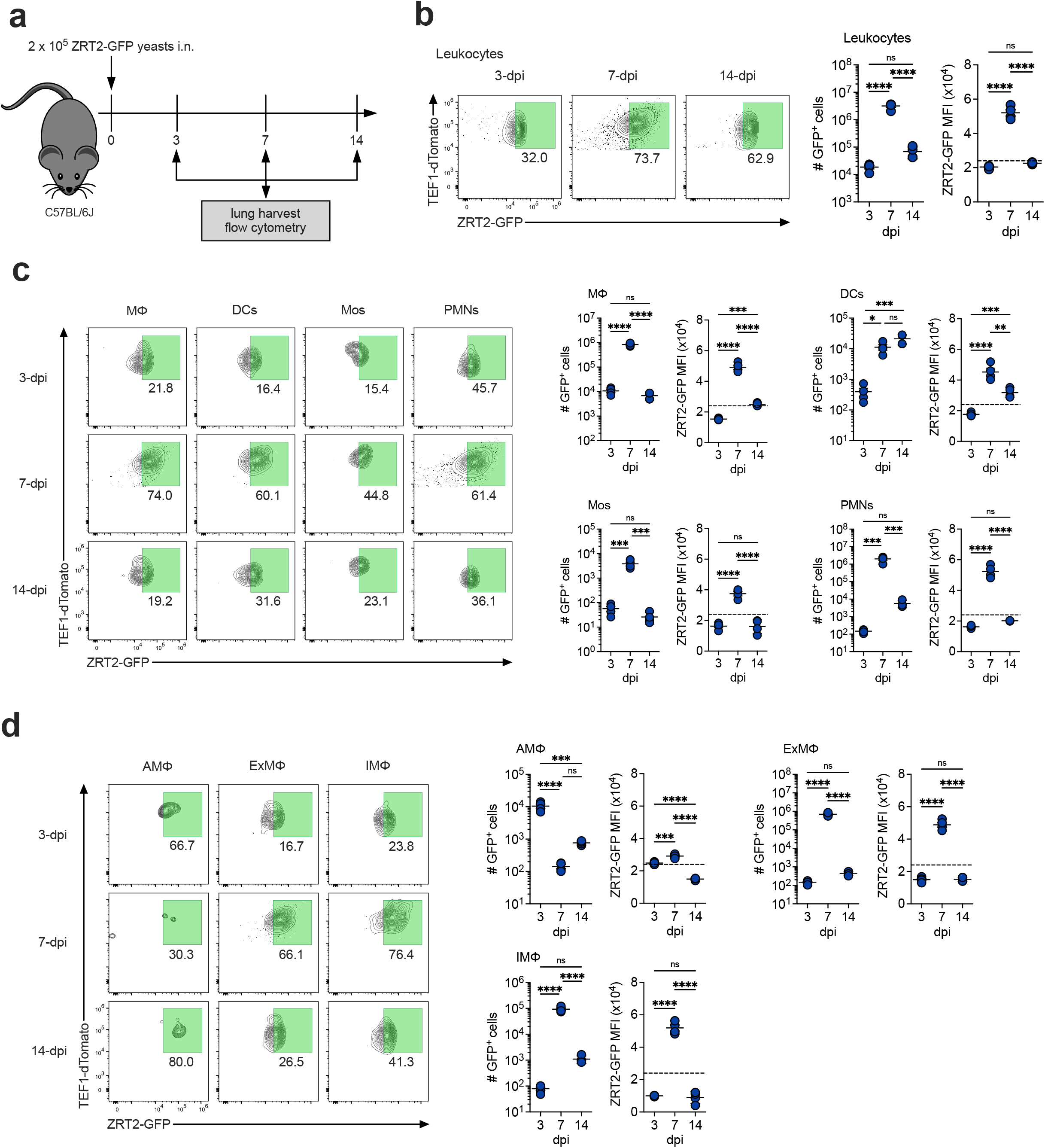
ZRT2-GFP expression peaks at 7-dpi in lung immune cells. (**a**) Experimental diagram. ZRT2-GFP expression and MFI at 3-, 7-, and 14-dpi in (**b**) total leukocytes, (**c**) MΦ, DCs, Mos, and PMNs, and (**d**) AMΦ, ExMΦ, and IMΦ from mouse lungs as measured by spectral flow cytometry. Dashed line represents the ZRT2-GFP cutoff equal to 4 standard deviations from ZRT2-GFP MFI at 2-hpi.

Analysis of lung leukocytes revealed maximal ZRT2-GFP expression at 7-dpi (Fig. 3b), congruent with previous work demonstrating ZRT2 is required for survival during adaptive immunity (25). We next optimized a 14-parameter spectral flow cytometry panel (Fig. S4) to interrogate ZRT2 expression in lung MΦ, DCs, monocytes (Mos), neutrophils (PMNs), and MΦ subpopulations – alveolar MΦ (AMΦ), interstitial MΦ (IMΦ), and exudate MΦ (ExMΦ) (30). MΦ represented the highest proportion of ZRT2-GFP^+^ immune cells at 3-dpi, whereas PMNs and MΦ contained the most ZRT2-GFP^+^ cells at 7-dpi (Fig. 3c). At 14-dpi there are equal numbers of ZRT2-GFP^+^ MΦ, DCs, and PMNs (Fig. 3c). In MΦ, at 3-dpi the predominant ZRT2-GFP^+^ cells were AMΦ, representing 56% of the total. ExMΦ and IMΦ were the major ZRT2-GFP^+^ MΦ populations at 7-dpi (Fig. 3d). At 14dpi, IMΦ and ExMΦ comprised 73% of total ZRT2-GFP^+^ cells (Fig. 3d). ZRT2-GFP^+^ AMΦ decrease between 3-7dpi but reappear between 7-14dpi (Fig. 3d). This loss may be attributable to cell death (31). These data strongly suggest that Zn deprivation is maximal at the onset of adaptive immunity.

### Anti-GM-CSF augments ZRT2-GFP expression in vivo

We inquired whether GM-CSF regulates ZRT2-GFP expression in vivo. We blocked endogenous GM-CSF and determined fungal burden and reporter expression at 3- and 7-dpi (Fig. 4a). Anti-GM-CSF increased CFUs at 7-dpi, but not 3-dpi (Fig. 4b). At 7-dpi, ZRT2-GFP expression was increased in total lung leukocytes from GM-CSF-neutralized mice (Fig. 4c). No changes in ZRT2-GFP expression were detected at 3-dpi in lung immune cells or MΦ (Fig. 4d and 4e). At 7-dpi, GFP MFI was heightened in MΦ, DCs, and PMNs from anti-GM-CSF-treated infected animals, while no changes in reporter expression and MFI between control and anti-GM-CSF mice were observed in Mos (Fig. 4f). ExMΦ represented most of the enhanced ZRT2-GFP expression observed in lung MΦ from recipients of anti-GM-CSF at 7dpi (Fig. 4g). These data imply that abrogation of GM-CSF signaling results in a more Zn-deprived environment within yeast-containing phagosomes, and this finding was quite contrary to our expectations.

**Figure 4.**
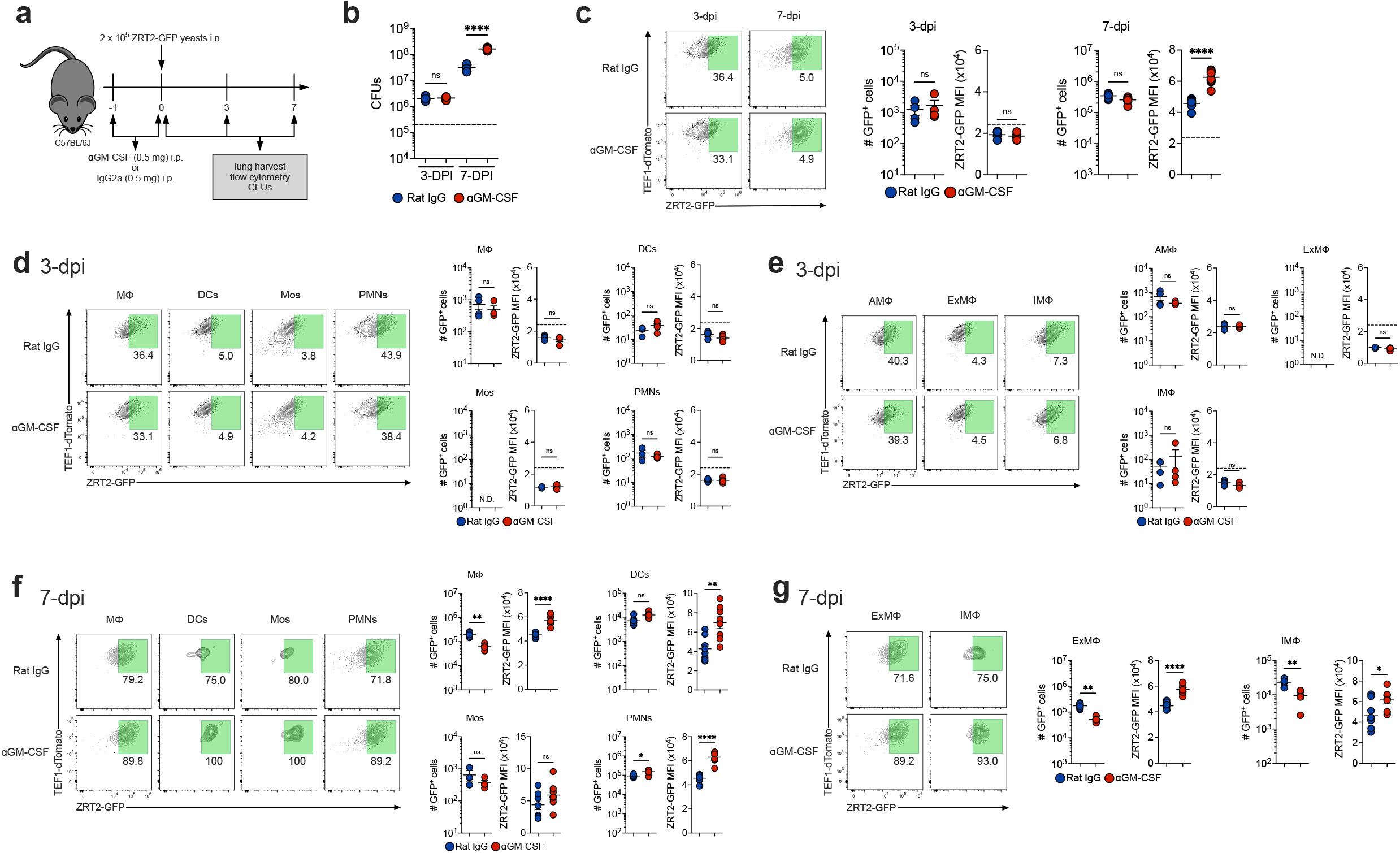
GM-CSF neutralization exhibited no effects at 3-dpi and increases ZRT2-GFP expression at 7-dpi in lung immune cells. (**a**) Experimental design. (**b**) CFUs at 3-dpi and 7-dpi. (**c**) ZRT2-GFP expression and MFI from rat IgG control (blue) and anti-GM-CSF (red) leukocytes at 3-dpi and 7-dpi. ZRT2-GFP expression and MFI from 3-dpi Rat IgG control (blue) and anti-GM-CSF (red) in (**d**) MΦ, DCs, Mos, and PMNs at (**e**) AMΦ, ExMΦ, and IMΦ. ZRT2-GFP expression and MFI from 7-dpi Rat IgG control (blue) and anti-GM-CSF (red) in (**f**) MΦ, DCs, Mos, and PMNs at (**g**) AMΦ, ExMΦ, and IMΦ. All data were measured by spectral flow cytometry. Dashed line represents the ZRT2-GFP cutoff equal to 4 standard deviations from ZRT2-GFP MFI at 2-hpi.

As a control, we neutralized IFNγ and probed ZRT2-GFP expression at 7-dpi (Fig. 5a). Increased ZRT2-GFP^+^ leukocytes were present in anti-IFNγ-treated lungs, but GFP MFI was similar between groups (Fig. 5b). We observed increased numbers of ZRT2-GFP^+^ DCs, Mos, and PMNs in anti-IFNγ-treated lungs, but no changes in GFP MFI among these populations (Fig. 5c). IMΦ exhibited a higher ZRT2-GFP expression and increased GFP MFI (Fig. 5d).

**Figure 5.**
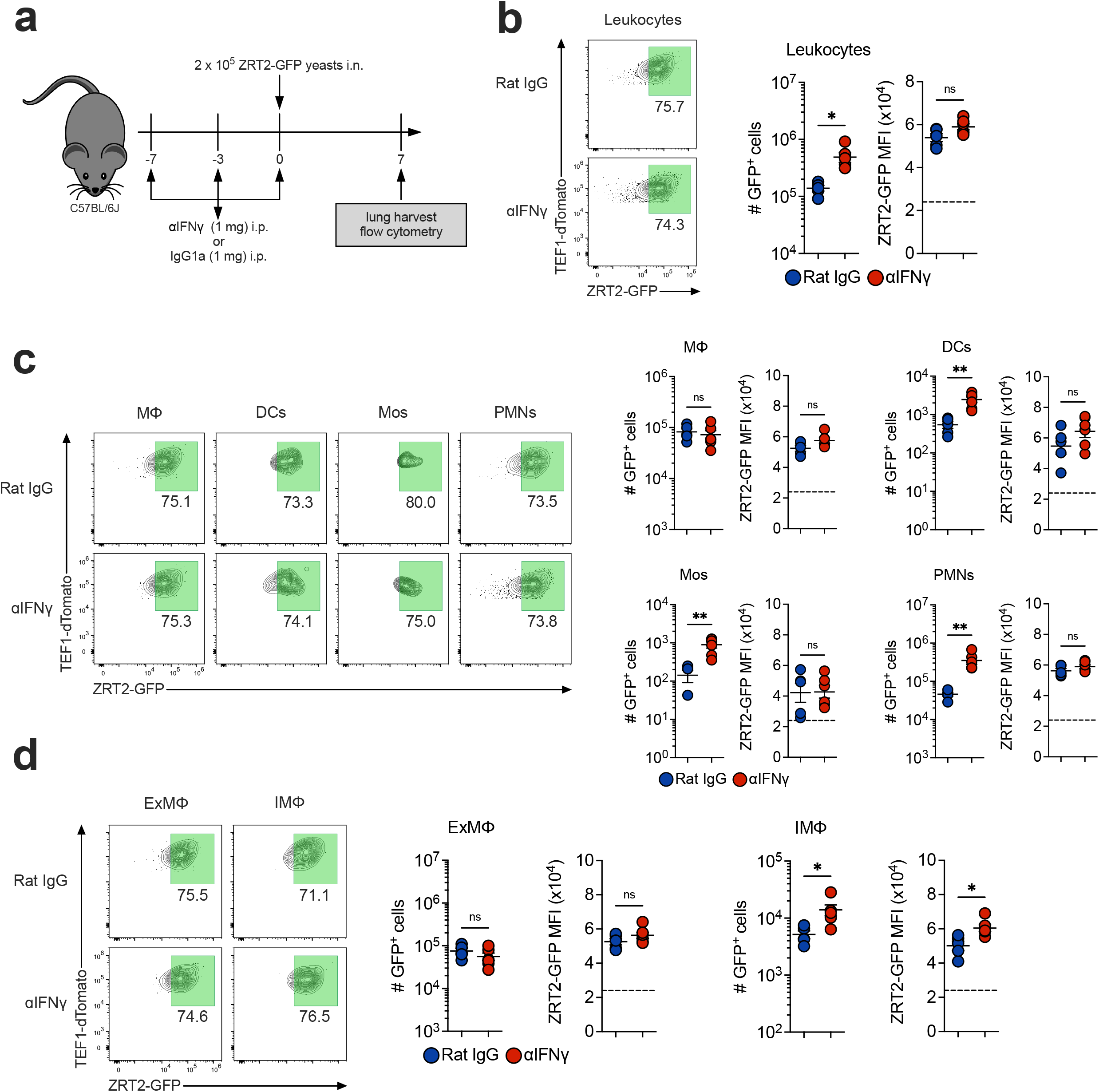
IFNγ neutralization does not affect ZRT2-GFP expression at 7-dpi in lung immune cells. (**a**) Experimental design. ZRT2-GFP expression and MFI from rat IgG control (blue) and anti-IFNγ (red) mice in (**b**) total leukocytes, (**c**) MΦ, DCs, Mos, and PMNs, and (**d**) AMΦ, ExMΦ, and IMΦ from mouse lungs as measured by spectral flow cytometry. Dashed line represents the ZRT2-GFP cutoff equal to 4 standard deviations from ZRT2-GFP MFI at 2-hpi.

### ZRT2-GFP expression is altered in *Csf2^-/-^*and *Mt1/2^-/-^* mice at 7-dpi

We complemented these studies by examining GM-CSF knockout (*Csf2^-/-^*) mice (Fig. 6a). The lungs of infected *Csf2^-/-^* mice contained considerably more CFUs (Fig. 6b). ZRT2-GFP^+^ lung leukocytes were similar at 7-dpi from *Csf2^-/-^* mice compared to WT; however, the GFP MFI in leukocytes from *Csf2^-/-^* mice was increased (Fig. 6c), mirroring results in anti-GM-CSF treated mice. GFP MFI was elevated in *Csf2^-/-^* MΦ, DCs, and PMNs, but not Mos (Fig. 6d). While there were fewer ZRT2-GFP^+^ IMΦ and ExMΦ, both populations manifested increased GFP MFI (Fig. 6e). The finding that MΦ enhance ZRT2 when GM-CSF is absent suggests another factor(s) drives expression.

**Figure 6.**
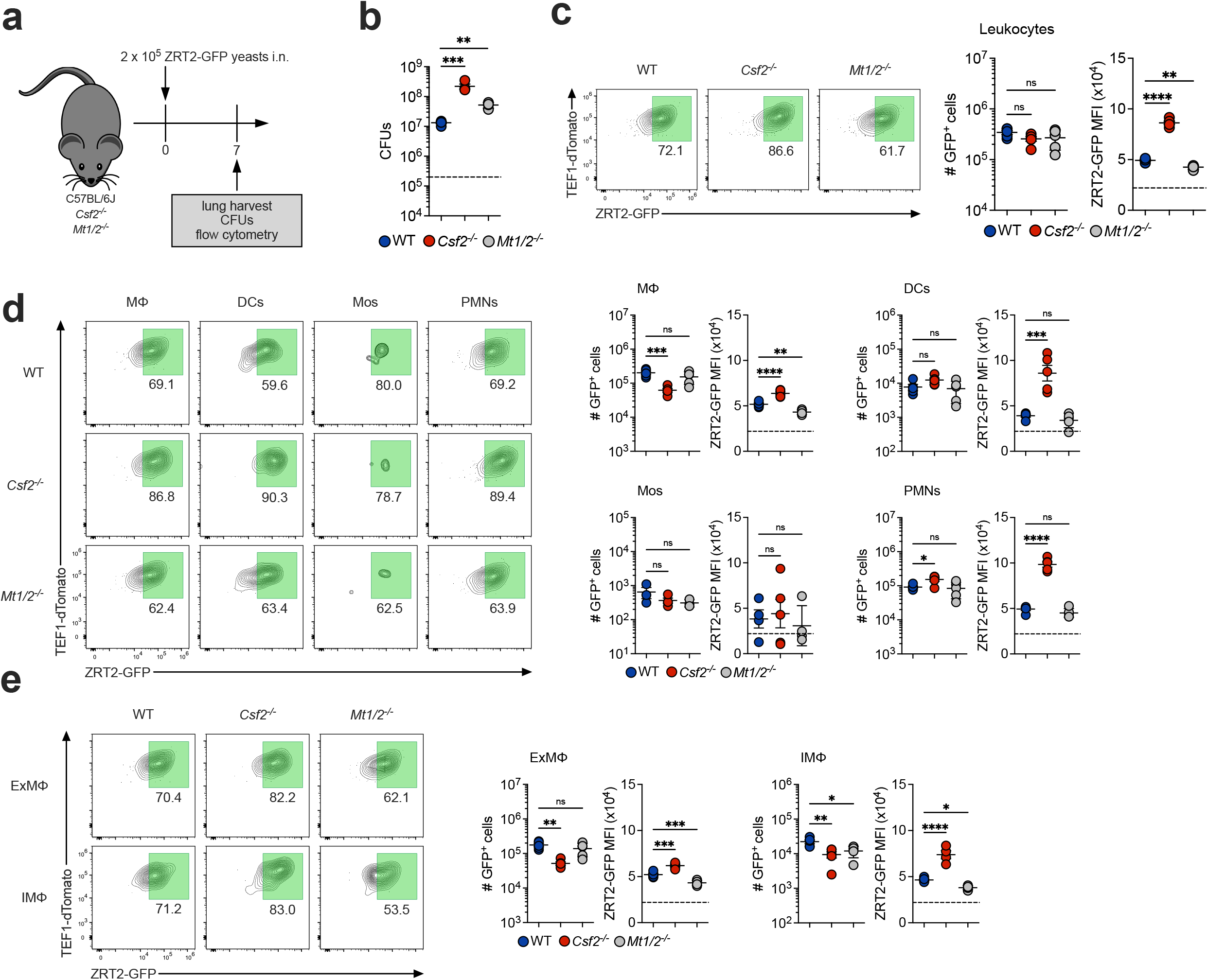
*Csf2^-/-^*mice exhibit increased ZRT2-GFP expression and ZRT2-GFP MFI is reversed in *Mt1/2^-/-^*mice compared to WT controls at 7 dpi. (**a**) Experimental diagram. (**b**) CFUs. ZRT2-GFP expression and MFI from WT (blue), *Csf2^-/-^* (red), and *Mt1/2^-/-^* (grey) mice in (**b**) total leukocytes, (**c**) MΦ, DCs, Mos, and PMNs, and (**d**) ExMΦ and IMΦ from mouse lungs as measured by spectral flow cytometry. Dashed line in (**b**) represents original inoculum. Dashed line in (**c**), (**d**), and (**e**) represents the ZRT2-GFP cutoff equal to 4 standard deviations from ZRT2-GFP MFI at 2-hpi.

We were perplexed by the above results and queried if any gene deletion in mice would trigger upregulation of ZRT2 in infected lung immune cells. We probed expression of this transporter in *Mt1/2^-/-^*mice. We selected these animals since they lack the major intracellular storage proteins for Zn. GFP MFI was decreased in leukocytes from *Mt1/2^-/-^* mice at 7-dpi (Fig. 6c). The number of ZRT2-GFP^+^ cells did not differ between groups in any of the four immune cell populations. MΦ from *Mt1/2^-/-^* mice exhibited decreased ZRT2-GFP MFI (Fig. 6d) as did IMΦ and ExMΦ (Fig. 6e). These data confirm the role of MTs 1 & 2 in Zn regulation in MΦ and support the fidelity of the reporter.

### M-CSF regulates ZRT2 expression when GM-CSF is absent

Since M-CSF activates the antifungal machinery of MΦ, we hypothesized it may contribute to ZRT2 expression (26). We first determined if M-CSF-stimulated BMDMs altered ZRT2-GFP expression. We exposed reporter-infected BMDMs to vehicle, GM-CSF, M-CSF, or IFNγ and assessed ZRT2-GFP. Treatment of BMDMs with GM-CSF or M-CSF induced a 60% increase in ZRT2-GFP^+^ cells and a 3-fold increase in GFP MFI (Fig. 7a). Addition of 100 µM Zn diminished GFP MFI in all groups (Fig. 7a). Thus, M-CSF establishes a Zn-deprived phagosomal environment.

**Figure 7.**
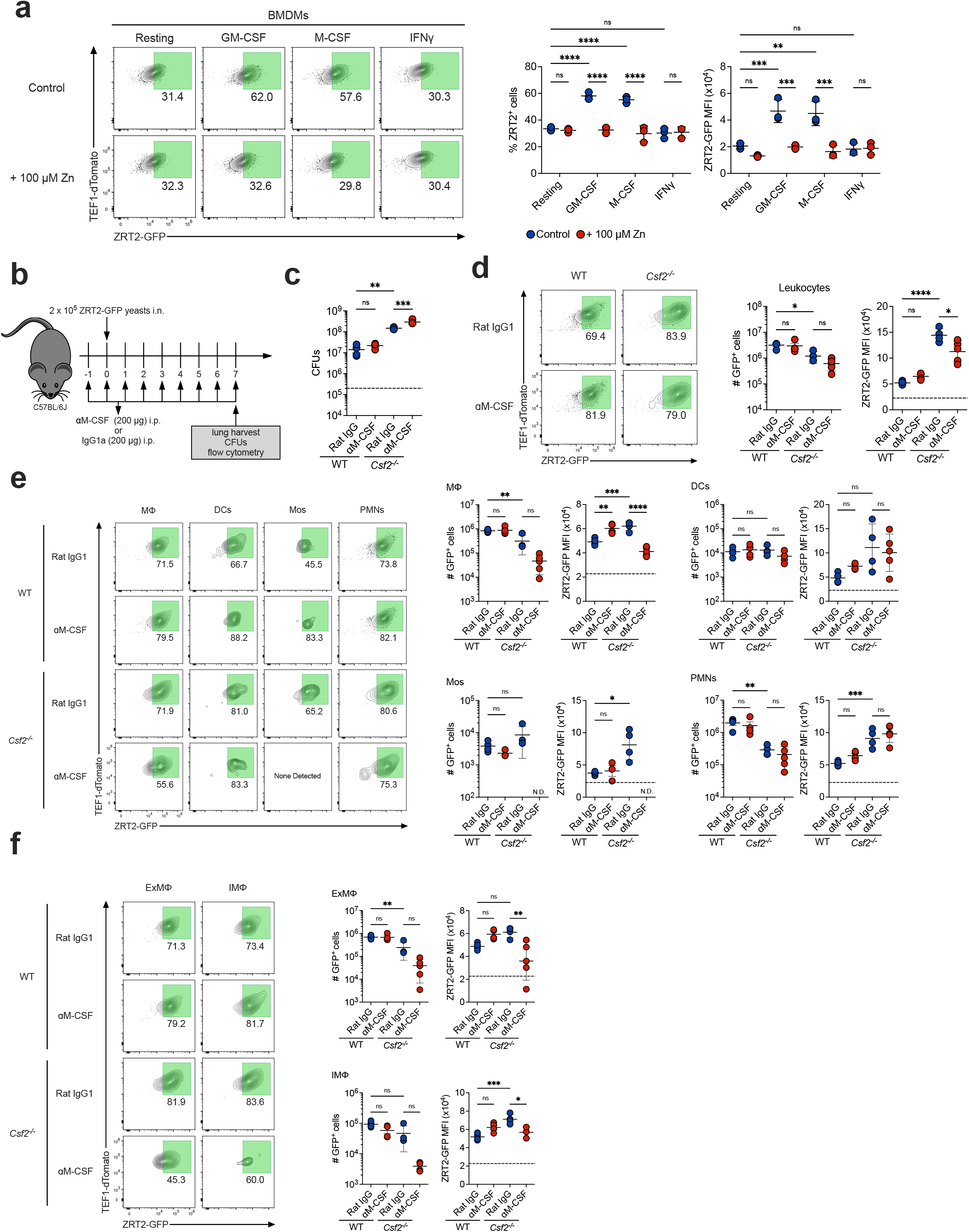
Ablation of both GM-CSF and M-CSF reverses ZRT2-GFP expression at 7-dpi. (**a**) ZRT2-GFP % and GFP MFI from ZRT2-GFP-infected BMDMs unstimulated, or stimulated with GM-CSF, M-CSF, or IFNγ 48-hpi in RPMI only (control) or Zn replete (+ 100 µM Zn) conditions. (**b**) Experimental design. (**c**) CFUs at 7-dpi from WT and *Csf2^-/-^* mice treated with Rat IgG isotype control (blue) or anti-M-CSF (red). ZRT2-GFP expression and MFI from WT and *Csf2^-/-^* mice treated with rat IgG isotype control (blue) or anti-M-CSF in (**d**) total leukocytes, (**e**) MΦ, DCs, Mos, and PMNs, and (**f**) ExMΦ and IMΦ from mouse lungs as measured by spectral flow cytometry. Dashed line in (**c**) represents original inoculum. Dashed line in (**d**), (**e**), and (**f**) represents the ZRT2-GFP cutoff equal to 4 standard deviations from ZRT2-GFP MFI at 2-hpi.

We next investigated if M-CSF contributed to ZRT2-GFP expression in vivo. We ablated M-CSF signaling by treating WT or *Csf2^-/-^* mice with anti-M-CSF mAb or isotype control and determined fungal burden and reporter expression at 7-dpi (Fig 7b). Lung CFUs were similar between WT given control antibody or anti-M-CSF; however, anti-M-CSF treatment increased fungal burden in *Csf2^-/-^* mice (Fig. 7c). While leukocytes from anti-M-CSF-treated mice exhibited no differences in GFP MFI compared to controls in WT mice, anti-M-CSF treatment reversed the enhanced GFP MFI in *Csf2^-/-^* mice (Fig. 7d). This change was specific for lung MΦ from *Csf2^-/-^*mice (Fig. 7e). ZRT2-GFP expression was reduced in ExMΦ and IMΦ from doubly-deficient mice (Fig. 7f). The increase in ZRT2- GFP expression in mice lacking GM-CSF is in part due to compensation by M-CSF.

## Discussion

Zinc is an essential divalent metal for cellular function, and its import by microbes, including *H. capsulatum*, during host residence is paramount for viability and virulence (8, 32, 33). In this study, we developed a ZRT2-GFP Zn reporter that specifically sensed and responded to changes in Zn abundance using 2 approaches: 1) supplementation of chelated medium with low and high concentrations of Zn, and 2) chelation of Zn by TPEN followed by re-addition of Zn. Both approaches upregulated the ZRT2-GFP reporter by 60 – 80%. The former approach enabled us to conclude that the Zn reporter unequivocally responds to changes in Zn availability, since ICP-MS analysis showed Zn was the only metal altered in yeast cells recovered from BMDMs in low Zn medium. Fungal orthologues of this importer manifest different specificities in transport of divalent cations. The *Cryptococcus gattii* ZRT2 orthologue is repressed by Fe (34). Conversely, ZRT2 orthologues in other fungal species only import Zn (32, 35–37). Although we reported that *H. capsulatum* ZRT2 expression is slightly repressed by Fe supplementation (25), our recent data substantiate that Zn is the sole metal that controls ZRT2 in *H. capsulatum*.

Reporter yeasts from GM-CSF-, but not IFNγ-, treated BMDMs exhibited a 3-fold increase in GFP MFI. These findings are not unexpected given that GM-CSF activation of BMDMs exclusively restricts Zn availability to *H. capsulatum*-containing phagosomes. Zn deprivation is linked with enhanced phagocyte-generated ROS to restrict fungal growth (22). Therefore, the possibility existed that ZRT2 responds to changes in ROS and Zn starvation. However, that was not the case since scavenging ROS did not alter ZRT2-GFP expression in GM-CSF-treated BMDMs.

Analysis of the evolution of ZRT2 in lung leukocytes revealed that increased expression was not observed until the onset of adaptive immunity, i.e., 7-dpi. By day 14, this expression had waned. Hence, the environmental cues that dictate the amount of labile Zn in *H. capsulatum*-containing phagosomes are not operational until the initiation of T cell-mediated immunity and are extinguished as clearance begins. The peak of ZRT2-GFP expression is tightly associated with the impact that GM-CSF has on host control of infection. Its absence only impacted fungal burden coincident with adaptive immunity.

We hypothesized that the lack of GM-CSF tempers ZRT2 expression in lung MΦ. By eliminating GM-CSF, the postulate was that the removal of a key element triggering Zn deprivation decreases ZRT2 expression. While in vivo disruption of GM-CSF predictably caused a loss of host control of infection, it unexpectedly resulted in a 4-6-fold increase in ZRT2-GFP expression. These data are counter to our in vitro data whereby GM-CSF treatment of BMDMs elicited an increase in ZRT2-GFP. One explanation to account for these discrepant results is that it is difficult to recreate the phagolysosomal environment of in vivo isolated cells and account for the plethora of signals that a lung phagocyte confronts. Nevertheless, the data indicate that other mechanisms maintain Zn stress within infected phagosomes in the absence of GM-CSF.

Given the surprising in vivo finding, we sought a model of infection to affirm that the ZRT2 reporter was not increased in any transgenic mouse. We capitalized on our development of *Mt1/2^-/-^* mice that lack the two major Zn-binding proteins downstream of GM-CSF activation. Analysis of the Zn reporter in lungs of mice devoid of MTs 1&2 revealed a blunted ZRT2-GFP expression. This result confirmed that the ZRT2 reporter behaved faithfully in vivo. The data also imply that loss of MTs does not initiate a compensatory mechanism to drive ZRT2-GFP expression and Zn deprivation.

One explanation of these discrepant results is contribution of M-CSF, another cytokine shown to inhibit *H. capsulatum* (26). A study with a mouse osteoblast line established a connection between Zn homeostasis and M-CSF (38). Hence, we explored the possibility that M-CSF influences ZRT2 expression in WT or GM-CSF deficient mice. While anti-M-CSF did not alter fungal burden or ZRT2-GFP expression in WT mice, it increased fungal burden in GM-CSF-deficient mice and conversely reduced ZRT2 expression comparable to that of WT mice. This outcome strongly suggests that M-CSF in the absence of GM-CSF contributes to phagosomal Zn restriction. Yet, the alleviation of Zn restriction in anti-M-CSF-treated *Csf2^-/-^*mice was dissociated from the elevated CFUs in doubly-deficient mice.

Investigating fungal sensing of Zn availability using fluorescent reporters provides unique insights into the phagolysosomal environment. Little is known about which populations of lung MΦ offer *H. capsulatum* yeasts a permissive replicative environment versus a growth-retarding environment. Dual RNA-sequencing of *M. tuberculosis*-infected MΦ revealed that AMΦ allow growth whereas IMΦ restrict growth (39–41). We showed that at 3dpi the majority of *H. capsulatum* yeasts resided in AMΦ and at 7dpi yeasts were within IMΦ and ExMΦ, suggesting a shift in *H. capsulatum* residence coincident with adaptive immunity. These two populations were most prominent in detecting altered expression of ZRT2. Thus, they are the primary target of mediators from T cells that calibrate host defenses.

Our study has limitations. First, we did not examine the role of ZRT1 that may act in concert with ZRT2. Second, we did not directly measure Zn in yeast cells from infected mice since that would have required lungs from many mice to capture yeast cells in sufficient quantities within MΦ subpopulations. Another confounding factor is that not all yeast cells upregulate ZRT2. This may be a matter of the dynamic nature of gene expression or may indicate that not all MΦ respond similarly or simultaneously to the environmental signals that drive increased expression of ZRT2.

In summary, we have developed a fluorescent reporter to assay changes in Zn homeostasis in mouse lungs. A few studies have probed intraphagosomal metal sensing using this approach (21, 35). Herein, we highlighted the complex nature of Zn regulation in *H. capsulatum*-infected phagocytes and provided a tool to probe various immune cell populations in host lungs over the course of infection. This study serves as a springboard to promote contemporary studies into the complex relationship between intracellular pathogens and the mammalian host.

## Materials and Methods

### Mice

Four to six-week-old male WT C57BL/6 mice and *Csf2^-/-^*mice were purchased from The Jackson Laboratory (Bar Harbor, ME). The original *Mt1/2^-/-^* strain (129S7/SvEvBrd-*Mt1^tm1Bri^Mt2^tm1Bri^*/J) was backcrossed > 9 times to C57BL/6J mice and genetic analysis revealed them to be > 99% C57BL/6J. All animals were housed under specific pathogen-free conditions in the University of Cincinnati’s Department of Laboratory Animal Medical Services facility which is accredited by the Association for Assessment and Accreditation of Laboratory Animal Care. All experiments were performed in accordance with the Animal Welfare Act guidelines of the National Institutes of Health.

### *H. capsulatum* strains and growth conditions

*H. capsulatum* strains used in this study (Table S1) were derived from the G217B clinical isolate. Yeast cells were maintained in *Histoplasma* macrophage medium (HMM) (4 µM Zn) with or without supplementation of FeSO4, ZnSO4, or CuSO4, where described (42). Uracil-auxotrophic *H. capsulatum* strains were supplemented with 100 µg/mL uracil. Yeasts were grown at 37°C at 200 rpm for 3 days. For growth on solid medium, 0.75% w/v agarose was added to HMM and supplemented with 25 µM FeSO4. For Zn deficient conditions, yeasts were grown in chelated HMM, and Cu^2+^, Fe^2+^, Mg^2+^, Ca^2+^, and Mn^2+^ were restored to their original quantity, as determined by ICP-MS (22).

### Generation of the *Histoplasma* ZRT2-GFP zinc reporter

To generate the ZRT2 promoter-GFP fusion, 1829 base pairs (bp) upstream of the ZRT2 coding sequence (pZRT2) were PCR amplified (primers: P1, P2; Table S2) from genomic DNA using primers designed with *Avr II* and *Asc I* restriction enzyme sites. The fragment was cloned into pCR™2.1 Vector (Invitrogen) to generate pLB110 and transformed into *E. coli* DH5ɑ (Invitrogen). pLB110 plasmid DNA was extracted using the Zippy Plasmid Mini Prep Kit (Zymo Research) and digested with *Avr II* and *Asc I*. This fragment was subcloned into pCR623, a *URA5*+ T-DNA vector, between the *Avr II* and *Asc I* restriction sites directly upstream of the coding sequence of GFP (pZRT2-*gfp*) and transformed into *E. coli* DH5ɑ. The sequence of ZRT2-GFP, pLB112, was confirmed by Sanger sequencing (University of Maine DNA Sequencing Facility, Orono, ME). pLB112 was inserted by *Agrobacterium tumefaciens*-mediated integration into the genome of *Histoplasma* strain OSU233 (43). *A. tumefaciens* bacteria containing pLB112 were cocultured with OSU233 *H. capsulatum* yeasts atop a sterile Whatman #5 filter on solid induction medium (0.5% D-glucose; 40 mM MES, pH 5.3; 350 µM cystine; and 5 mM PO4) containing 0.1 mM acetosyringone (ThermoFisher) for 48 hours at 26°C. The filter was transferred to solid HMM plates devoid of uracil to select for transformants, and 200 µM cefotaxime to counter select *A. tumefaciens*. The plates were incubated at 37°C for until transformants appeared. Transformants were inoculated into liquid HMM devoid of uracil and incubated for 5–7 days at 37°C. Genomic DNA was extracted from each transformant and PCR-verified (Primers: P3, P4, P5; (Table S2) to contain the integrated ZRT2-GFP construct. Positive transformants were probed for dTomato and GFP expression by incubation in Zn-free HMM for 24 hours in clear-bottom, black microtiter plates (ThermoFisher) and measurement of GFP (excitation 485 nm/ emission 528 nm) and dTomato (excitation 555 nm/ emission 585 nm) fluorescence using a BioTek Synergy plate reader (BioTek, Winooski, VT). Transformants with at least 3-fold increase in GFP expression were selected as candidate Zn reporter strains.

### Generation and infection of BMDMs with *H. capsulatum*

Bone marrow from femurs and tibias of 8–12-week-old mice was isolated and differentiated in complete RPMI-1640 (C-RPMI) (Cytiva) containing 10% fetal bovine serum (FBS), 55 µM 2-mercaptoethanol (Sigma), and 10 µg/mL gentamicin sulfate (Sigma) supplemented with 10 ng/mL recombinant mouse GM-CSF (Biolegend). Cells were incubated at 37°C with 5% CO2. GM-CSF was added every 3 days. After 7 days, non-adherent cells were removed, and adherent cells incubated with trypsin/EDTA (Corning) and gently scraped with a cell scraper. BMDMs were seeded into 24-well tissue culture plates (Corning) at 10^6^ cells/well and rested overnight at 37°C. BMDMs were stimulated for 2 hours prior to infection with vehicle or 10 ng/mL of each cytokine: GM-CSF (Biolegend), M-CSF (Miltenyi), or IFNγ (Biolegend). Yeasts were diluted in C-RPMI and BMDMs infected at the indicated multiplicity of infection (MOI). Two hours post-infection, plates were washed 3 times with warm C-RPMI to remove non-phagocytized yeasts. Where indicated, cells were exposed to 10 µM TPEN (Cayman Chemical Company), 100 µM ZnSO4 (Sigma), 25 µM FeSO4 (Sigma), or 1 µM CuSO4 (Sigma). To inhibit ROS generation, BMDMs were treated with 200 µM apocynin (Sigma) 24 hours before and at the time of infection.

### Gene Expression Analysis

*H. capsulatum* RNA was isolated by mechanical disruption with 0.5 mm glass beads, extraction with TRIzol™ (Invitrogen), and precipitated with ethanol, as previously described (44, 45). Genomic DNA was removed via DNase digestion and column purification using the PureLink™ RNA Mini Kit (Invitrogen). Complimentary DNA was prepared from 1 µg of RNA using a Reverse Transcription Systems Kit (Promega). Quantitative PCR (qPCR) was performed on an ABI Prism 7500 (Applied Biosystems) using SYBR green (ThermoFisher). Primer pairs were designed to yield an approximate 200 bp product (Table S3). Oligonucleotides were synthesized by Integrated DNA Technologies, Inc. (IDTDNA). Transcripts were normalized to the glyceraldehyde 3-phosphate dehydrogenase (*GAPDH*) gene. Gene expression was determined using the *ΔΔ*Ct method (46) and normalized to control conditions.

### Infection of mice and treatment with monoclonal antibodies (mAb)

Six to 8-week mice were anesthetized via isoflurane inhalation and intranasally (i.n.) infected with 2 x 10^5^ *H. capsulatum* yeasts. To determine baseline ZRT2-GFP expression, single cell suspensions from mice infected for 2 hours were analyzed by flow cytometry. In some experiments, mice were given neutralizing mAb to cytokines or isotype control antibodies. Mice were injected intraperitoneally (i.p.) with 500 µg of control mAb rat IgG2a (Leinco Technologies, Inc.) or 500 µg MP1-22E9 mAb anti-GM-CSF (Leinco Technologies, Inc.) on -1- and 0dpi (47). To neutralize IFNγ, mice were injected i.p. with 300 µg of clone XMG 1.2 mAb anti-IFNγ (Cell Culture Company) or 300 µg isotype control mAb rat IgG1a (BioXCell) on -7dpi, -3dpi, and 0dpi (48). To neutralize M-CSF, mice were injected i.p. with 200 µg of isotype control mAb rat IgG1a (BioXCell), or 200 µg of 5A1 mAb to M-CSF (Bio X-cell) daily on days -1- to 6dpi. To measure fungal burden, lungs were homogenized in HMM, serially diluted, and plated on solid HMM at 37°C for 7–14 days, or until colonies appeared.

### Isolation of single cells from lungs

Lungs were homogenized in a GentleMACs™ Dissociator (Miltenyi) containing 5 mL dissociation buffer (RPMI-1640 (Cytiva) with 25 mM HEPES and 10 µg/mL gentamicin sulfate) supplemented with 2 mg/mL Collagenase-D (Roche) and 100 U DNaseI (Roche). The samples were incubated on a 200-rpm shaker at 37°C for 30 minutes. The digested lung cells were percolated through a 70 µm cell strainer and washed with HBSS (Corning) containing 2 mM EDTA. Red blood cells were lysed using ACK lysis buffer and resuspended in PBS containing 2% FBS, 2 mM EDTA, and 2 mM NaN3. Viable cells were enumerated by an Automated Cell Counter (BioRad).

### Flow Cytometry

One million lung cells were stained. Fc receptors were blocked for 10 minutes with CD32/CD16 mAb (Leinco Technologies, Inc.). Cells were incubated with Zombie UV™ Fixable Viability Dye (BioLegend) and stained with a mixture of fluorochrome-conjugated antibodies (Table S4). Samples were fixed with 2% v/v paraformaldehyde (ThermoFisher Scientific). All antibodies were titrated to optimal concentration before experiments were performed. Data were obtained on a Cytek™ Aurora Spectral Analyzer (Cytek) using SpectroFlo^®^ software and analyzed using FlowJo™ *v10.9* (Tree Star). UltraComp eBeads™ (Invitrogen) and ArC™ Amine Reactive Compensation Beads (Life Technologies) were used for all single-color controls. Gating strategies are shown in Fig. S4 and determined using unstained and fluorescent minus-one samples. All flow cytometric data were acquired using equipment maintained by the Research Flow Cytometry Core (RFCC) in the Division of Rheumatology at Cincinnati Children’s Hospital Medical Center.

### ICP-MS

After treatment as described in each experiment, Zn reporter infected BMDMs were lysed with 0.1% SDS in water for 30 minutes on ice. Lysates were centrifuged and the *H. capsulatum* pellets were analyzed by ICP-MS. Fungal pellets were rinsed 3 times with cold PBS and transferred to a metal-free vial. Total metal analysis was performed by ICP-MS after acid mineralization. In brief, 100 µL of 1:1 trace grade nitric acid/ doubly deionized water and 20 µL of 500 ppb internal standard mixture were added to the pellet and heated on a dry bath for 30 minutes at 60°C. The vials were vented, and the temperature increased to 95°C for another 30 minutes. Samples were then cooled to room temperature and brought to a final volume of 1 mL with double deionized water.

Metal quantitation was performed on an Agilent 7500ce ICP-MS system (Agilent Technologies) with a Scott double-pass spray chamber and MicroMist nebulizer (Glass Expansion), a standard 2.5 mL torch, and nickel cones. The system was operated in collision mode with 3.5 mL/min of He and a calibration range from 0.2 to 25 ppb with the external calibration method. ^66^Zn, ^63^Cu, ^59^Co, ^56^Fe, ^31^P, ^34^S, and ^55^Mn were quantified with ^45^Sc, ^89^Y, and ^115^In as internal standards. Sulfur was used as an internal mass index as reported previously (49).

### Statistical Analysis

All data are represented as individual data points and the mean ± standard deviation (SD). Data were analyzed by Student’s *t-*test, one-way and two-way ANOVA, followed by Tukey’s multiple comparisons test or Sidak’s multiple comparisons test. analysis was performed using Prism *v9.5.1* (GraphPad Software). Significant differences are indicated in graphs with asterisk symbols (*, *P* < 0.05; **, *P* < 0.01; ***, *P* < 0.001, ****, P < 0.0001) or no significant differences (n.s.).

## Acknowledgments

The authors thank Dr. Chad Rappleye (The Ohio State University) for supplying the OSU233, WU15, and OSU326 *H. capsulatum* strains, and the backbone TEF1-GFP *A. tumefaciens* vector (pCR623), Daniela Amado, Kamila Cuervo, and Katie Morrice, for technical assistance and Alyssa Sproles for help with flow cytometry. The work was supported by NIH grants AI106269, AI133797, and a University of Cincinnati Dean’s fund grant to George Deepe. The Cytek Aurora analyzer within RFCC was purchased with NIH S10OD025045.

## Supplemental Material

**Figure S1.**
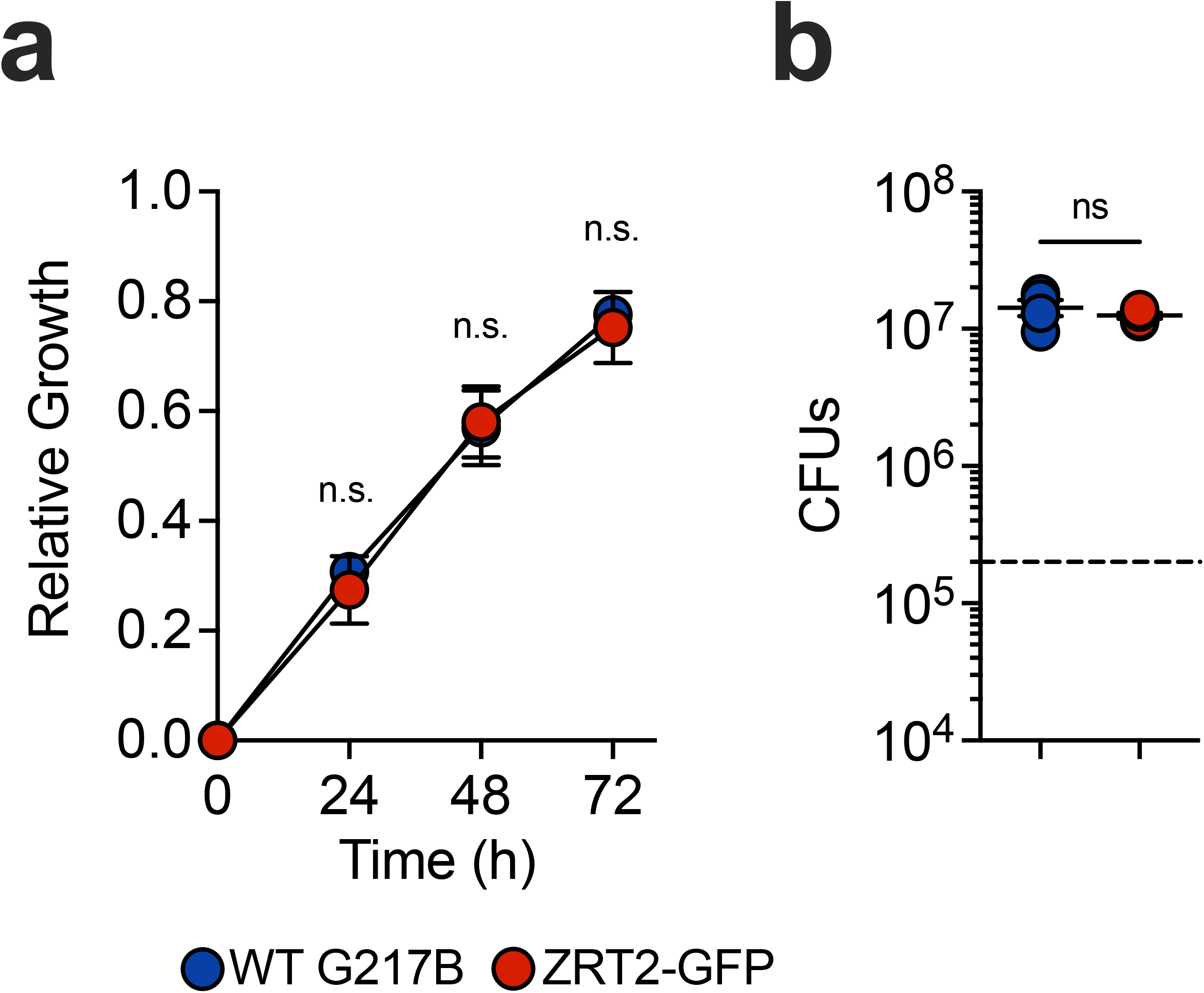
The Zn reporter strain exhibits no defects in growth or virulence. (**a**) Growth of WT and ZRT2-GFP *H. capsulatum* yeasts in liquid HMM. (**b**) CFUs or ZRT2-GFP *H. capsulatum* yeasts recovered from mouse lungs 7-days post-infection.

**Figure S2.**
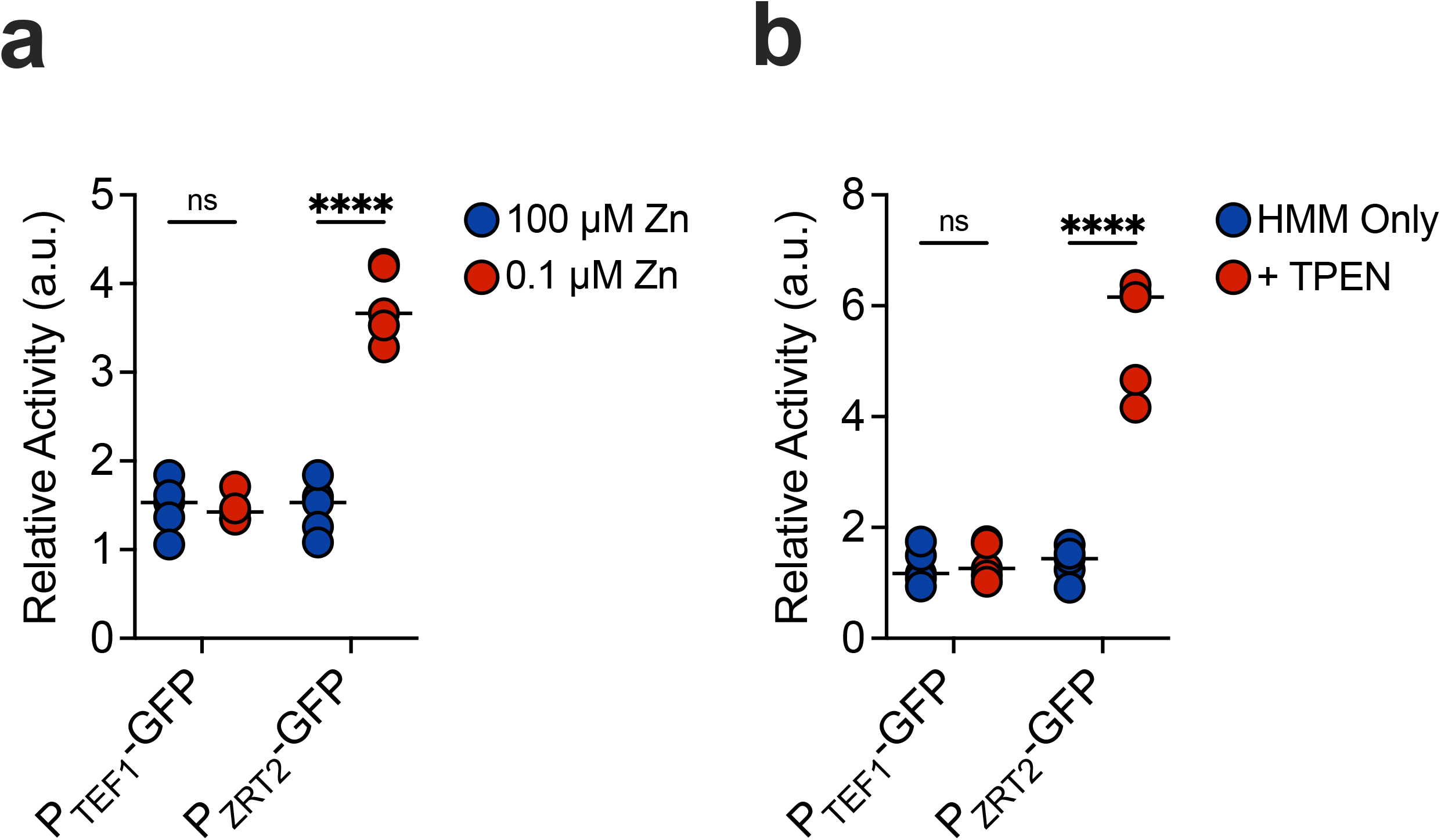
The Zn reporter strain responds to Zn-limited growth conditions. (**a**) Relative activity of ZRT2-GFP or TEF1-GFP *H. capsulatum* yeasts in chelated liquid HMM supplemented with 100 µM ZnSO_4_ or 0.1 ZnSO_4_, respectively. (**b**) Relative activity of ZRT2-GFP or TEF1-GFP *H. capsulatum* yeasts in liquid HMM treated with 10 µM TPEN or vehicle. dTomato and GFP fluorescence was measured 48 hours after the start of experiments.

**Figure S3.**
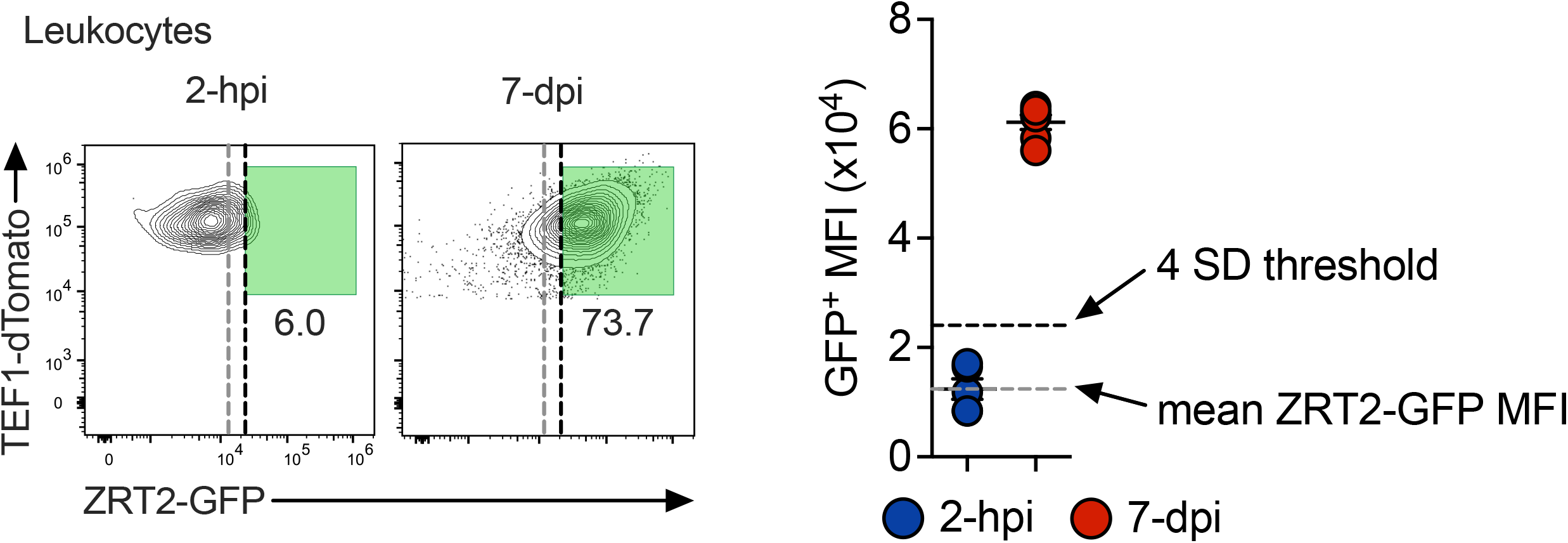
Setting ZRT2-GFP threshold for in vivo measurement of reporter activity. WT mice were infected with 2 x 10^5^ ZRT2-GFP *H. capsulatum* yeasts i.n. and sacrificed at 2-hours post-infection (2-hpi) and 7-days post-infection (7-dpi) and ZRT2-GFP MFI was calculated. 4 standard deviations (SD) from the mean MFI at 2-hpi was determined as the threshold for the ZRT2-GFP reporter to be “on” vs. “off.”

**Figure S4.**
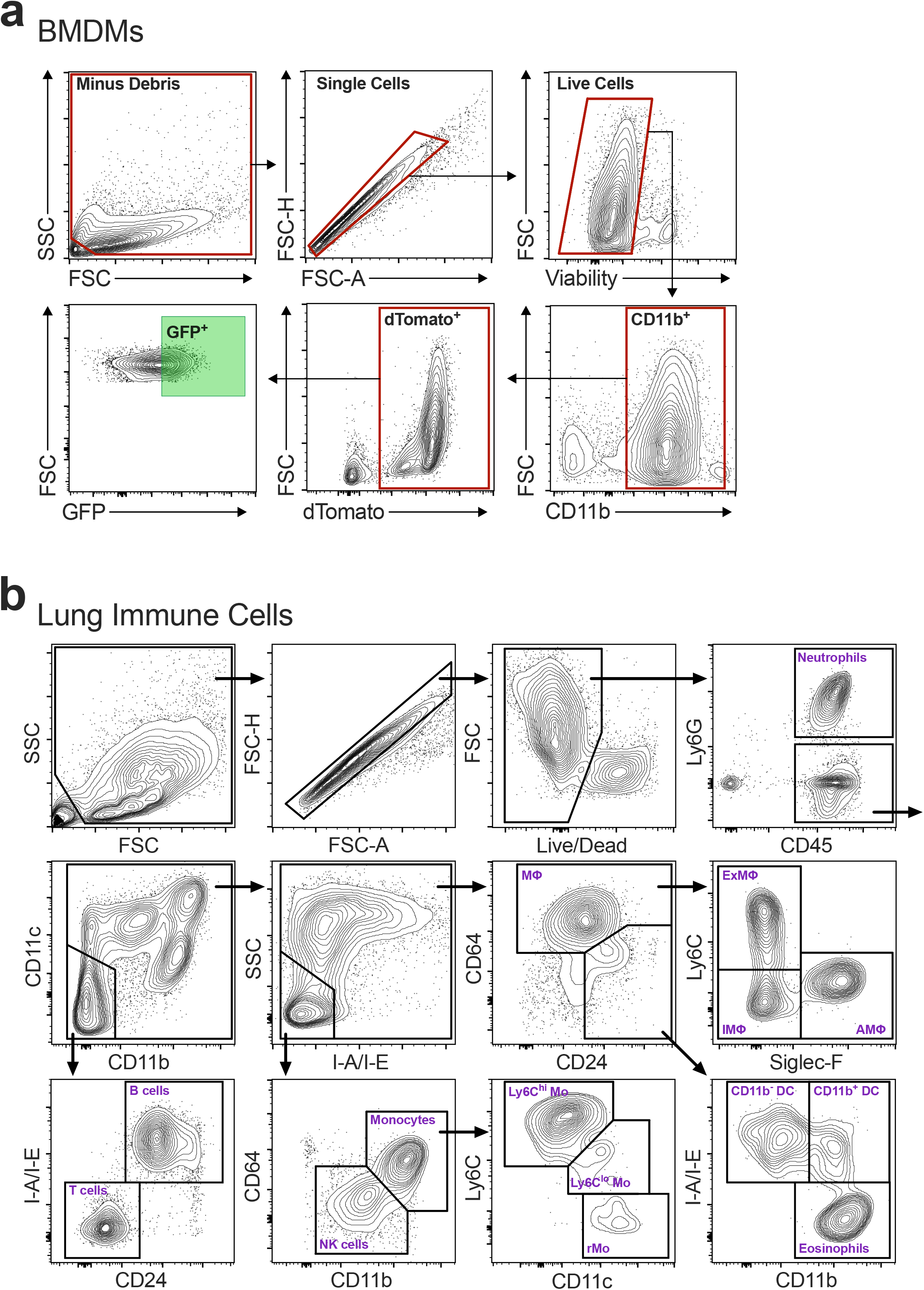
Spectral Flow Cytometry Gating Strategies. Gating strategies for (**a**) BMDMs and (**b**) lung immune cell populations. Data was acquired on a Cytek Aurora Spectral Cytometer.

**Table S1:** *H. capsulatum* Strains.

| strain | Genotype<sup>1</sup> |
| --- | --- |
| G217B | Wild type NAm2 isolate (ATCC 26032) |
| WU15 | *ura5-42Δ* |
| OSU233<sup>3</sup> | *ura5-42Δ* zzz:pQS01 (*apt3*, *P<sub>TEF1</sub>-rfp*) |
| OSU326<sup>3</sup> | *ura5-42Δ* zzz:pCR623 (*URA5*, *P<sub>TEF1</sub>-gfp*) |
| UC25 | *ura5-42Δ* zzz:pQS01 (*apt3*, *P<sub>TEF1</sub>-rfp*) zzz:pLB111 (*URA5*, *P<sub>ZRT2</sub>-gfp*) |

<sup>1</sup> gene designations:
zzz:: plasmid integration at an undetermined chromosomal location
*apt3*: aminoglycoside phosphotransferase (G418 resistance)
*gfp*: green-fluorescence protein (*EGFP*)
*rfp*: red-fluorescence protein (tdTomato)
*URA5*: orotate phosphoribosyltransferase
*TEF1*: translation elongation factor EF-1 $\alpha$
*ZRT2*: high affinity zinc transporter
<sup>2</sup>source: Shen Q, et al., 2018

**Table S2:**
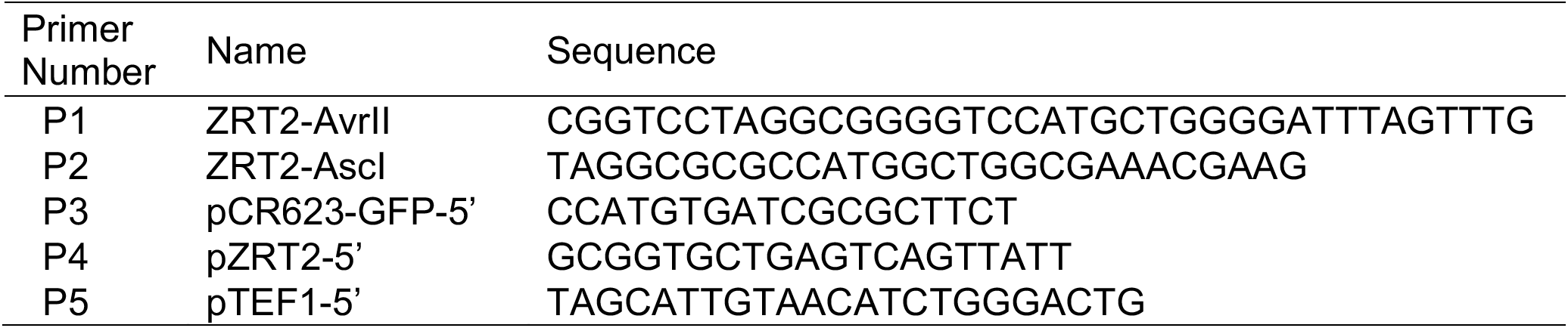
PCR Primers Used in This Study.

**Table S3:**
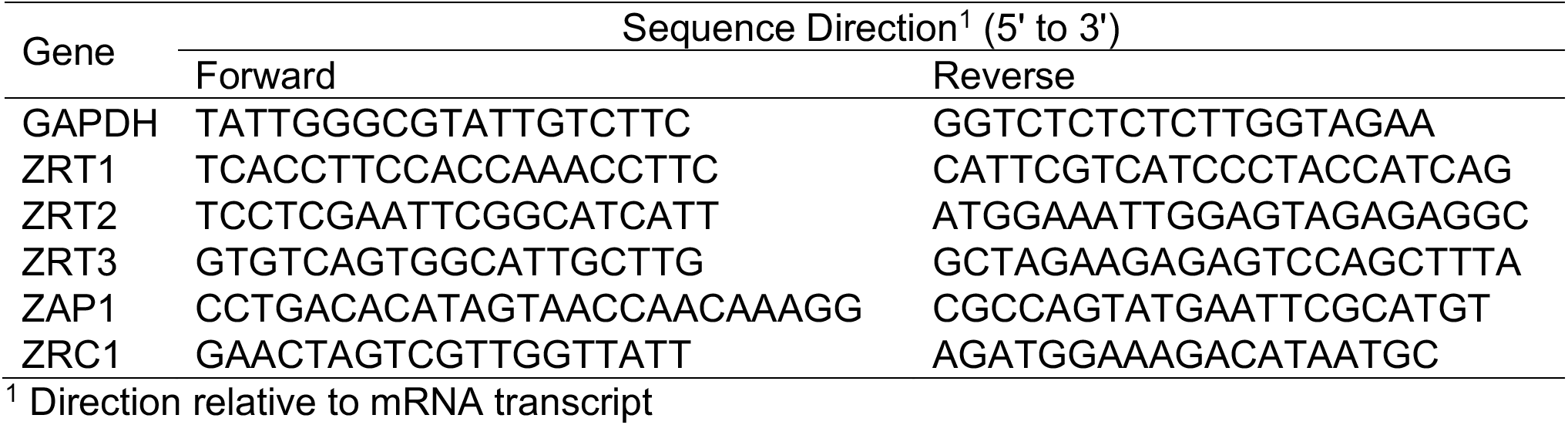
qPCR Primers Used in This Study.

**Table S4:**
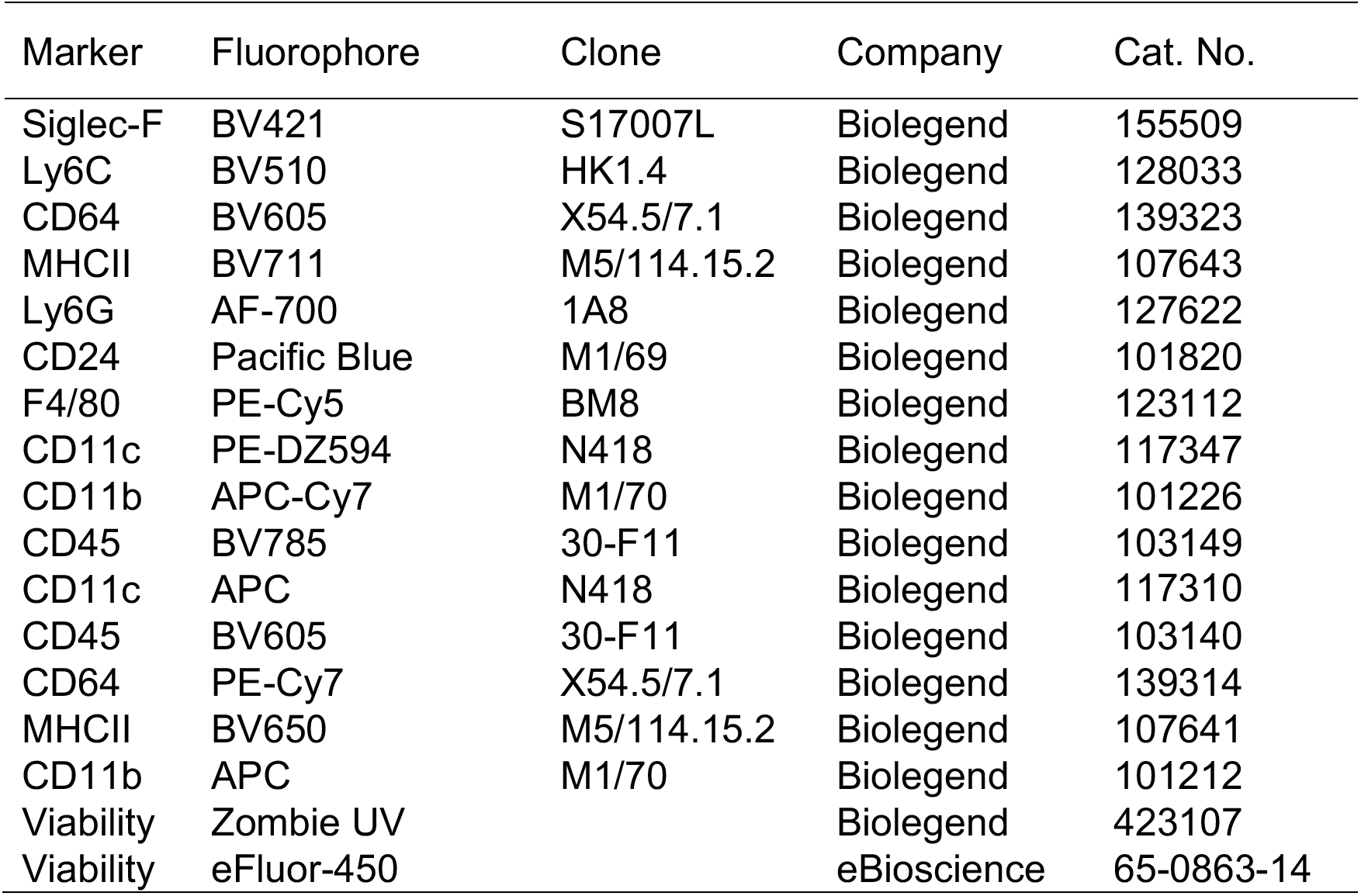
Flow cytometry antibodies.

